# Microplastic Exposure Promotes Amyloid Misfolding and Metabolic Impairment at Sub-Lethal Doses in an In Vitro Cellular Model of Alzheimer’s Disease

**DOI:** 10.64898/2025.12.22.696023

**Authors:** Iran Augusto Neves da Silva, Agnes Paulus, Valeriia Skoryk, Kar Yan Su, Fátima Herranz-Trillo, Oxana Klementieva

## Abstract

Microplastics and nanoplastics (MNPs) are ubiquitous environmental pollutants with increasing implications for human health. While their presence in human tissues is established, the molecular mechanisms driving their potential neurotoxicity remain unclear. This study investigates the impact of polystyrene (PS) on amyloid protein misfolding and cellular metabolism using Optical Photothermal Infrared (O-PTIR) spectroscopy, a label-free, sub-diffraction imaging technique. Our results reveal that PS exposure promotes pathological protein misfolding, specifically increasing β-sheet-rich conformations, and disrupts metabolic homeostasis at sub-lethal doses. These suggest that the microplastic surface acts as a catalytic scaffold for amyloid aggregation, driving cellular dysfunction prior to acute toxicity. This identifies a plausible molecular pathway by which environmental MNP pollution contributes to the risk and progression of neurodegenerative diseases, highlighting the need for risk assessments that look beyond simple cell survival.

## INTRODUCTION

Microplastics (MPs, <5mm) and nanoplastics (NPs, <100 nm), collectively referred to as MNPs, are pervasive pollutants found in air, soil, food packages, and water [1, 2]. Polystyrene (PS), specifically, is a critical priority for toxicological assessment, as it is one of the six most abundant plastic polymers, accounting for a significant portion of global plastic production [3].

Human exposure to MNPs is pervasive and occurs through multiple pathways, including ingestion, inhalation, and dermal absorption [4–6]. Sources include everyday items like food containers, which unintentionally release MPs into the body. High-temperature exposure, such as when drinking hot beverages, can significantly increase the release of billions of microparticles and nanoparticles from sources like plastic teabags and paper cups [7]. This high environmental mobility and bioaccessibility has led to general health concerns, including endocrine disruption, reproductive toxicity, and cancer [6, 8].

MNPs can cross biological barriers, potentially accumulating in the brain what may lead to cognitive issues [1]. Recently, MNPs were detected to bioaccumulate in human brain tissue [9]. Additionally, as reported by Leslie et al. (2024), who showed not only the presence of plastic particles in postmortem human brain samples but also a correlation between polymer exposure and molecular changes in the brain associated with neurodegenerative disease. Their findings suggest that plastic accumulation may modulate neuroinflammatory pathways and protein misfolding processes relevant to Alzheimer’s disease (AD) [10].

AD is a progressive neurodegenerative disorder characterized by memory loss, cognitive decline, and neuronal death. It is pathologically defined by the accumulation of extracellular amyloid plaques and intracellular neurofibrillary tangles [11]. While the exact cause of AD remains multifactorial, the potential for environmental pollutants to trigger surface-catalysed amyloid aggregation is a growing concern. Research indicates that polystyrene can modulate the formation of amyloid fibrils, with studies showing that particles around 400 nm significantly promote the primary nucleation step in proteins like hen egg-white lysozyme (HEWL) [2]. This effect is attributed to hydrophobic interactions occurring at the interface and junction of the protein corona that forms around the polystyrene particles [12, 13].

Notably, mechanistic studies utilizing elevated exposure models have successfully demonstrated that MNPs can induce oxidative injury and neuronal apoptosis in fetal thalamus and neuronal cell lines. However, the specific molecular role of the plastic surface in directly catalysing protein misfolding remains unexplored [14]. MNPs also interfere with lipid metabolism, potentially causing the accumulation of toxic lipid species in the brain [1, 2] and contributing to neuronal dysfunction [15–18]. Additionally, MNPs not only bind to lipids, altering their structure and function, but also induce broader cellular damage, including inflammation and cell death [19]. Despite growing concerns, the exact mechanisms by which MNPs contribute to AD processes remain poorly understood [20, 21].

In AD, Aβ peptides misfold and aggregate into a range of structural forms, from soluble oligomers to insoluble fibrils, which typically accumulate extracellularly to form amyloid plaques [22]. However, growing evidence indicates that intracellular aggregation and the structural conformation of Aβ particularly oligomeric β-sheet–rich species are more closely associated with neurotoxicity [23].

This study investigates whether microplastic exposure, specifically to PS, influences the intracellular aggregation of Aβ. We hypothesize that PS promotes conformational changes such as the formation of β-sheet–rich structures and oligomeric species that disrupt cellular homeostasis. To model this, we used N2A cells overexpressing human amyloid precursor protein (APP) carrying the Swedish mutation (N2Aswe).

To explore structural changes of Aβ(1-42), we employed optical photothermal infrared (O-PTIR) spectroscopy, which enables label-free, high-resolution structural imaging. O-PTIR combines the molecular specificity of infrared (IR) absorption with the spatial resolution of visible light, overcoming the diffraction limits of conventional IR techniques to achieve sub-micron resolution (~500 nm) [24]. The amide I region (1600–1700 cm^−1^), which reflects the C=O stretching vibrations of peptide bonds, is particularly informative for detecting structural transitions of proteins [25, 26].

Within the amide I region (1600–1700 cm^−1^), several sub-bands are particularly informative for amyloid research. β-sheet structures typically show strong absorption in the lower range (~1610–1640 cm^−1^), with aggregated or intermolecular β-sheets appearing around 1610–1625 cm^−1^. Native antiparallel β-sheets are observed near 1625– 1640 cm^−1^, while a high-frequency component around 1680–1695 cm^−1^ also corresponds to β-sheet structures. In contrast, α-helices and random coils appear at higher wavenumbers (~1640–1658 cm^−1^ and ~1640–1650 cm^−1^, respectively) and turns or loops are usually detected in the 1660–1680 cm^−1^ range. The ratio of these spectral features enables the discrimination of different amyloid conformations and aggregation states in situ [27, 28].

Our results provide direct mechanistic evidence that PS exposure drives pathological protein misfolding, specifically evidenced by a distinct shift toward β-sheet-rich conformations, with increased unordered and antiparallel β-sheet detected at ~1640 cm^−1^ and 1694 cm^−1^, respectively. Beyond structural alterations, PS exposure severely disrupted metabolic homeostasis, demonstrated by a substantial decrease in metabolic activity and lysosomal dysfunction. These critical findings, obtained through the sub-cellular resolution of O-PTIR, confirm that the microplastic surface acts as a catalytic scaffold for amyloid aggregation. This identifies a plausible molecular pathway by which environmental plastic pollution contributes to the etiology of neurodegenerative diseases, highlighting an urgent need to reassess the chronic health risks of MNP accumulation in the brain.

## MATERIALS AND METHODS

Commercially available labelled, 365/415 nm (DAPI) and 580/605 nm (CY5) PS beads FluoSpheres carboxylate-modified microspheres, with a diameter of 0.2 µm, were obtained from Thermo Fisher cat. F8805 and F8810 concentration of 2% of solids in 10 ml solution.

### Cell Culture and Exposure

N2A cells (ATCC CCL-131) were utilized in this study. Two variants were prepared: the N2A wild type (N2Awt), representing the non-transfected parental cell line, and the N2A cell line stably transfected with human APP carrying the Swedish mutation [29] (N2A APPSwe), which will be referred to as N2Aswe in further experiments. Both N2Awt and N2Aswe cells were cultured in a 1:1 ratio of DMEM and Opti-MEM (31985062; Gibco), supplemented with 10% FBS and 1% penicillin/streptomycin, and maintained at 37°C with 5% CO2. N2Aswe cells were selected for stable expression using 50 mg/ml Geneticin (10131027; Gibco) in their media. For experimental procedures, cells were exposed to PS concentrations of 0, 120, 240, and 480 µg/mL. These concentrations were selected to align with recent mechanistic investigations into PS-induced neurotoxicity and ROS-mediated injury, ensuring a comparable toxicological baseline for evaluating surface-catalysed protein misfolding[14]. and Aβ(1-42) for either 24 or 48 hours, once they reached 60% confluence.

### Metabolic Assessment of N2Aswe WST-1

Metabolic activity (i.e., cell viability) was assessed using the water-soluble and cell-permeable tetrazolium salt reagent (WST-1), which is reduced on the membrane of mitochondria in living cells [30]. Following intracellular cleavage of WST-1 by cellular mitochondrial dehydrogenases into formazan, the cell-permeable formazan is released into the cell culture media and can subsequently be measured by removing the liquid and reading it in a standard UV–vis plate reader. Absorbance can be viewed as proportional to the metabolic activity of living cells. Therefore, lower amounts of formazan indicate a reduced metabolic activity.

N2Aswe exposed to (0, 120, 240 and 480 µg/ml) during 24 and 48 hours, were incubated with 90 μL of complete medium and 10 μL of WST-1 (Roche, Sigma-Aldrich, USA) for 1 h at 37 °C in a humidified incubator with 5% CO2. Supernatant optical density was measured at 440 and 650 nm on an Epoch plate reader (Epoch plate reader, BioTek, Winooski, Vermont, United States) 440 nm was used as a representative for the specific conversion of WST-1 by metabolically active cells, and 650 nm wavelength was used as a reference wavelength for nonspecific absorbance. Complete medium without tissue served as a negative control. All assays were performed in independent biological triplicates (n=3).

### Lactate Dehydrogenase Assay

To assess the cytotoxic effect of Polystyrene (PS) and Aβ(1-42) exposure, a Lactate Dehydrogenase (LDH) cytotoxicity detection kit (ab197004, Abcam, Cambridge, UK) was used. Monomeric Aβ(1-42) was prepared from a stock solution of 250 mM in DMSO. The peptide was diluted in culture medium immediately prior to use to ensure it remained in its monomeric state for the duration of the 24-hour exposure. Both N2Awt and N2Aswe cells were cultured for 24 hours prior to exposure, with treatments conducted for an additional 24 hours. All conditions were performed in triplicate (n=3). The experimental groups were: PS Dose Response (0 (Untreated Control), 100, 120, 240, and 480 µg/mL PS), Abeta Alone (10 mM Aβ(1-42) in the media), and a Co-Exposure Group (480 µg/mL PS combined with 10 µM Abeta (1-42)). Following the 24-hour exposure, approximately 5 µL of supernatant from each treatment group were collected. The LDH Reaction Mix solution (95 µL) was prepared by combining 2 µL of Developer Mix I/LDH Substrate Mix, 4 µL of Pico Probe III/Pico Probe, and 89 µL of LDH Assay Buffer. 95 µL of the prepared LDH Reaction Mix solution was added to 5 µL of the collected supernatant in a 96-well plate. The plate was then incubated at room temperature, in the dark, for 10 minutes. Controls included: Background Control (Basal media only), Negative Control (Untreated cell supernatant), Lysate Control (Supernatant from Lysis Buffer-treated cells, representing maximum release), and a Positive Control (LDH positive control buffer provided by Abcam). Fluorescence was measured at an excitation/emission wavelength of 535 nm / 587 nm using the Cytation 5 Cell Imaging Multi-Mode Reader. Data is reported in Relative Fluorescence Units (RFU), calculated as per the manufacturer’s suggested protocol and normalized against the background media control. The percentage of cytotoxicity was calculated by normalizing the LDH release of the exposed samples against the maximum release (Lysate Control).

### Optical Photothermal Infrared Spectroscopy

For microspectroscopy we used mIRage system (Photothermal Spectroscopy Corp., Santa Barbara, CA, USA) at Lund University within the Integrated Vibration Spectroscopy-Microsom Laboratory for Molecular-Scale Biogeochemical Research. The IR source was a pulsed, tunable four-stage quantum cascade laser (QCL) operating in the IR with a repetition rate of 100 kHz, inducing localized heating and thermal expansion in the sample. This photothermal expansion in the sample was detected using a 532 nm green probe laser, enabling the acquisition of quantitative absorption spectra. Background spectra were collected on a built-in reference sample. For further details about the method can be found in the references [31–35]. For the experiment investigating Aβ fibrillization induced by PS, the following instrument settings were applied. Single spectra were collected within the 1300 cm^−1^ to 1800 cm^−1^ range using a standard silicon photodiode detector with a gain of 50x. The IR power was set to 100%, the probe power to 4.7 %, with an average of 10 spectra. Background spectra were obtained from a built-in reference standard to ensure consistency and calibration across measurements. The IR spectra were cut to the Amide I region, spanning 1500–1700 cm^−1^.

### Data analysis

The spectra obtained from the experiments were analysed using Quasar-M data mining software [36]. A linear baseline correction was applied to remove any systematic offset. The spectra were then vector-normalized to correct for variations in overall signal intensity. To enhance the number of discriminative spectral features, the Savitzky–Golay smoothing algorithm was used, implemented with a 13-point window and a second-order polynomial. To examine the impact of PS on N2Aswe cells, high-resolution hyperspectral maps were collected using an APD detector with a spatial resolution of 0.5 μm x 0.5 μm from individual cells (5 cells non-exposed and 6 cells exposed to PS) across the spectral range of 1000 cm^−1^ to 1750 cm^−1^. For each cell, between 370 - 400 spectra were acquired. The IR source power was set to 58%, the probe power to 0.5%, and 5 spectra were averaged per measurement. For the analysis, to focus on the region associated with protein secondary structure, the spectra were cut to the 1610–1710 cm^−1^ range (Amide I). To enhance spectral features and resolve overlapping spectral peaks relevant to structural protein analysis were improved by calculating the second-order derivative of each spectrum using the Savitzky–Golay algorithm with a 7-point window. To ensure data quality and robustness for all the collected spectra for the experiments, an unsupervised outlier detection method was applied. Approximately 10% of the spectra identified as anomalous were excluded using the LOF algorithm.

In order to analyse the intensity ratios, the total β-sheet content was determined by calculating the intensity ratio (Table 1) between 1630 cm^−1^ (corresponding to β-sheet structures) and 1658 cm-1 (to the total protein/ α-helices peak). Antiparallel β-sheet structures (1694 cm^−1^) were quantified by their intensity ratio relative to both total proteins (1658 cm^−1^) and total β-sheet structures (1630 cm^−1^). Similarly, unordered β-sheet structures (1640 cm^−1^) were calculated based on their intensity ratio relative to total proteins (1658 cm^−1^) and total β-sheet structures (1630 cm^−1^). Principal Component Analysis (PCA) was used to explore differences in the IR spectral profiles of N2Aswe cells exposed to PS compared to non-exposed controls. Prior to PCA, spectra were processed by calculating the second derivative, followed by baseline correction and normalization to the total protein peak at 1658 cm^−1^. PCA was performed on the 1610–1710 cm^−1^ region corresponding to Amide I vibrations. Components were retained based on the screen plot and cumulative explained variance, with the first four components accounting for 70% of the total variance. PCA scatter and loading plots were used to assess group separation and identify spectral features contributing to observed differences. All analyses were conducted on the pre-processed dataset, enabling reproducible and reliable comparisons across experimental conditions.

**Table 1.** Infrared Band Positions and Their Assignments to Biomolecular Vibrational Modes [27, 28, 37–40].

| Band position ($\text{cm}^{-1}$) | Vibrational Assignments, $\nu$ |
| --- | --- |
| 1602 | Polystyrene; aromatic ring; |
| 1630 | Major β-sheets; Amide I; |
| 1640 | Random coil/unordered structures; Amide I; |
| 1658 | α-helices in proteins; Amide I; |
| 1694 | Minor Antiparallel β-sheets; Amide I; |

$\nu$ – stretching mode

### Electron Microscopy

#### Transmission Electron Microscopy (TEM)

After 1 hour of aggregation, we placed a droplet of the PS + Aβ (1-42) mixture onto a 400-mesh, 3 mm copper grid with a carbon support film (EM Resolutions Ltd., Sheffield, UK) and let it incubate for 10 minutes. We then quickly washed the grid twice with Milli-Q water and left it to air dry until imaging. Imaging was performed on a JEOL 1400 PLUS (120kV).

#### Scanning Electron Microscopy (SEM)

For SEM analysis, individual samples of diluted PS, diluted Aβ (1-42), nd the combined PS + Aβ (1-42) mixture were prepared. Five microliters of each sample were deposited onto separate silicon wafers and air-dried completely. Prior to imaging, dried samples were sputter-coated with a thin layer of gold for enhanced conductivity. SEM imaging was carried out using a HITACHI SU3500 at the Microscopy Platform at the Department of Biology, with images acquired at an accelerating voltage of 5 kV and a working distance of 5470 µm.

### Fluorescent labelling

Amytracker^R^ 520 (Ebba Biotech, Solna, Sweden). According to the manufacturer’s protocol, Amytracker^R^ 520 was diluted from 1:1000 in the presence of DAPI diluted from 1:10000 and living tissue slices were incubated for 30 min at room temperature in a 24-well plate. The 82E1 primary antibody (IBL, 1:1000 dilution) was used to detect total Aβ, Ab-42 antibody conjugated with 555 Alexa (1:1000), while the OC78 (Abcam, 1:1000 dilution) primary antibody was applied to detect fibrillar oligomeric forms including soluble oligomeric forms. Cells were incubated with Alexa 568 goat anti-mouse and Alexa 648 goat anti-rabbit (dilution 1:1000) for 1 hour and then mounted with mounting media DAKO for observation. Primary antibodies used LAMP-1 (Abcam, 1:2000 dilution) as an indicator of lysosome formation, where PS were expected to accumulate, and anti-caspase-3 (Cell Signalling Technologies, 1:1000 dilution) as an apoptotic marker. (Amytracker 630 (Ebba Biotech, cat. number: A630-50EX, 1:1000 dilution) was also used to potentially detect amyloid accumulation. Cells were then incubated with Alexa 488 goat anti-rabbit (Invitrogen, cat. number: A11034) and Alexa 568 goat anti-rat (Invitrogen, cat. number: A11077) at a dilution of 1:1000. Imaging was done using BioTek Cytation 5 Cell Imaging Multi-Mode Reader and the epifluorescence Axio Zeiss microscope system. PBS-Tween20 was used as a washing buffer for the immunofluorescence, and 1% BSA in 1X PBST acted as both blocked buffer and antibody dilution buffer.

### Small-angle X-ray scattering (SAXS)

Small-angle X-ray scattering (SAXS) measurements were performed to determine the radius of gyration (Rg) of PS both alone and in the presence of (Aβ1–42) in physiological buffer at room temperature. All samples were carried out at CoSAXS beamline at the synchrotron facility MAXIV (Lund, Sweden), operated at a fixed energy (12.4 keV, l = 0.99 Å) with an incident X-ray beam focused on the detector. The sample-to-detector (Eiger2 X 4 M) distance was set at 10 m, yielding a q-range of 0.0033 < q < 0.26 Å-1, suitable for analysing PS diameter. The scattering intensity was recorded as a function of the scattering vector (q), and the data were processed to extract Rg values using the Guinier approximation [41, 42]. The final concentration of PS and Aβ1–42 in the mixtures was adjusted to 2 mg/ml and 0.04 mg/ml, correspondingly. The concentration of monomeric Aβ1–42 was set below detection limit to exclude contribution to scattering. Buffer-only controls were subtracted from all datasets to eliminate background contributions.

### Imaging

Imaging was acquired using the manual mode in a Cytation 5 multimode reader (Agilent Technologies, USA). Montage images were collected at 4× magnification. Bright field and DAPI imaging filter cube were used with the following acquisition settings: Bright: LED intensity: 10, integration time: 21, camera gain: 24, immunofluorescence: DAPI: LED intensity: 4, integration time: 5, camera gain: 1. GFP: LED intensity:3, integration time: 98 camera gain: 18. TEXAS: LED intensity: 6, integration time: 528 and 98 camera gain: 18. CY5 2, 1994 and 18, DAPI: LED intensity: 8, integration time: 5, camera gain: 1. Immunofluorescent images were acquired using the microscope Zeiss AxioImager (Zeiss Axio Imager M2) where images were collected at 40x magnification and used afterwards for the fluorescence intensity calculation of the specific antibodies mentioned above. At least eight images per group were collected. Quantitative analysis of the area and intensity of specific fluorescent signals was performed using FIJI software, with group comparisons made using the Mann–Whitney U-test.

## RESULTS

To determine if O-PTIR could effectively investigate PS in a label-free setting and under various conditions, we first evaluated its ability to analyse the spectral absorption characteristics of PS on CaF_2_ substrates. Our analysis identified distinct spectral markers at 1600 cm^−1^, 1494 cm^−1^, 1453 cm^−1^, and 1027 cm^−1^, for PS which could serve as references for future experiments. These markers are consistent with previous literature (**Figure 1**) [40, 43, 44].

**Figure 1.**
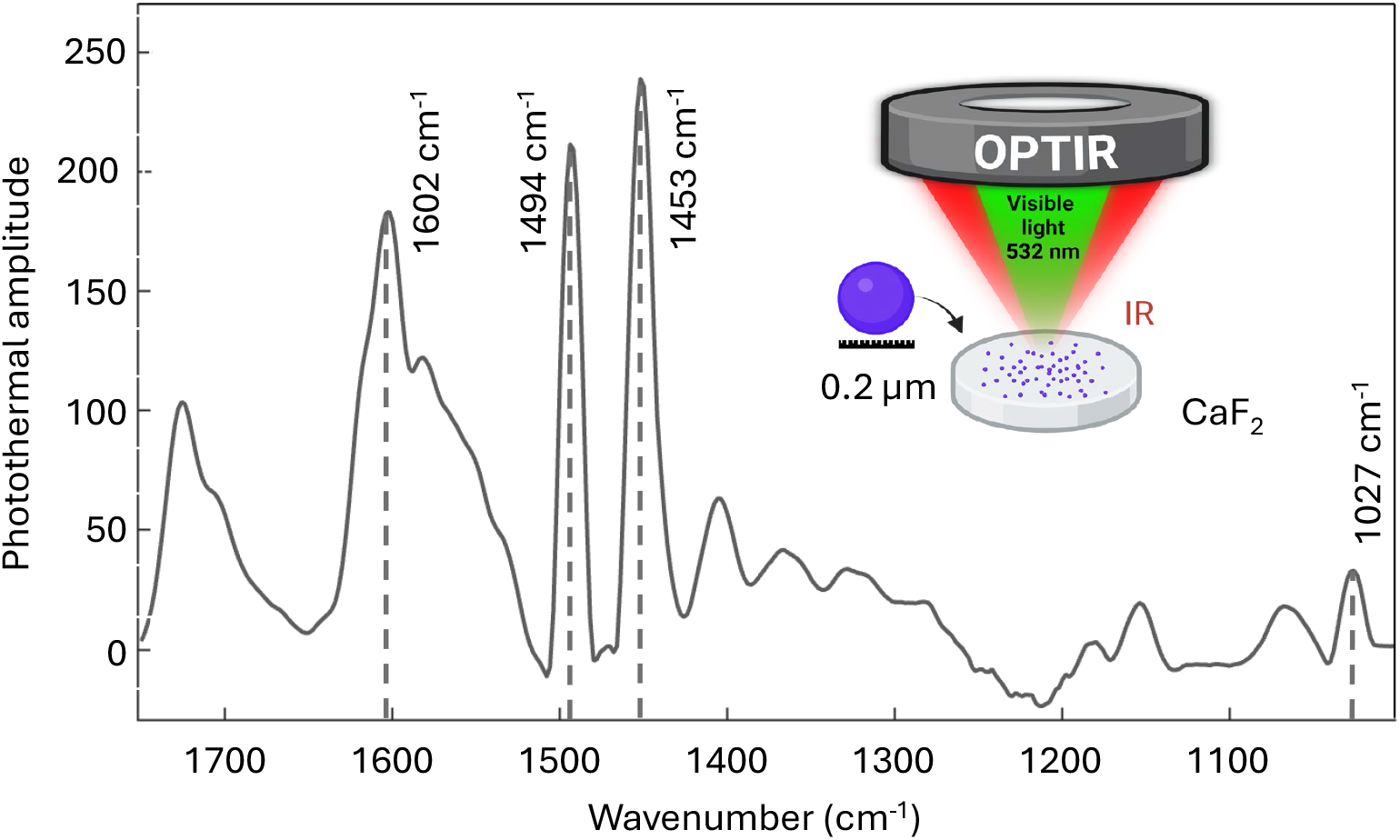
O-PTIR spectra of PS. The insert shows the principle of how O-PTIR operates: an IR laser is used to induce photothermal expansion within the sample, while green laser (532 nm) is used to probe the localized thermal effect. These specific absorption bands are consistent with previously reported literature [40, 43, 44]. O-PTIR schematics created in BioRender, Silva, I. (2025) https://BioRender.com/d94r687.

### Aβ fibrillization induced by the presence of PS

To investigate how PS microplastics influence amyloid protein structure, we incubated recombinant Aβ (1-42) with PS for 1 hour. (**Figure 2A**) provides a schematic overview of the experimental timeline, illustrating the separate observation of PS and Aβ, and their co-incubation over time. Fluorescence microscopy, presented in (**Figure 2B**), demonstrates the impact of PS on Aβ (1-42) aggregation. Initially, Aβ (1-42) alone appeared as smaller structures. At time zero (T0) of co-incubation, PS and Aβ (1-42) showed some initial aggregates, with a significant increase in aggregate size and fluorescent intensity observed after 1 hour of co-exposure. The accompanying graph in (**Figure 2B**.i) quantifies this observation, showing a significant increase in Aβ fluorescence size when co-incubated with PS, both at T0 and after 1 hour, compared to Aβ alone.

**Figure 2.**
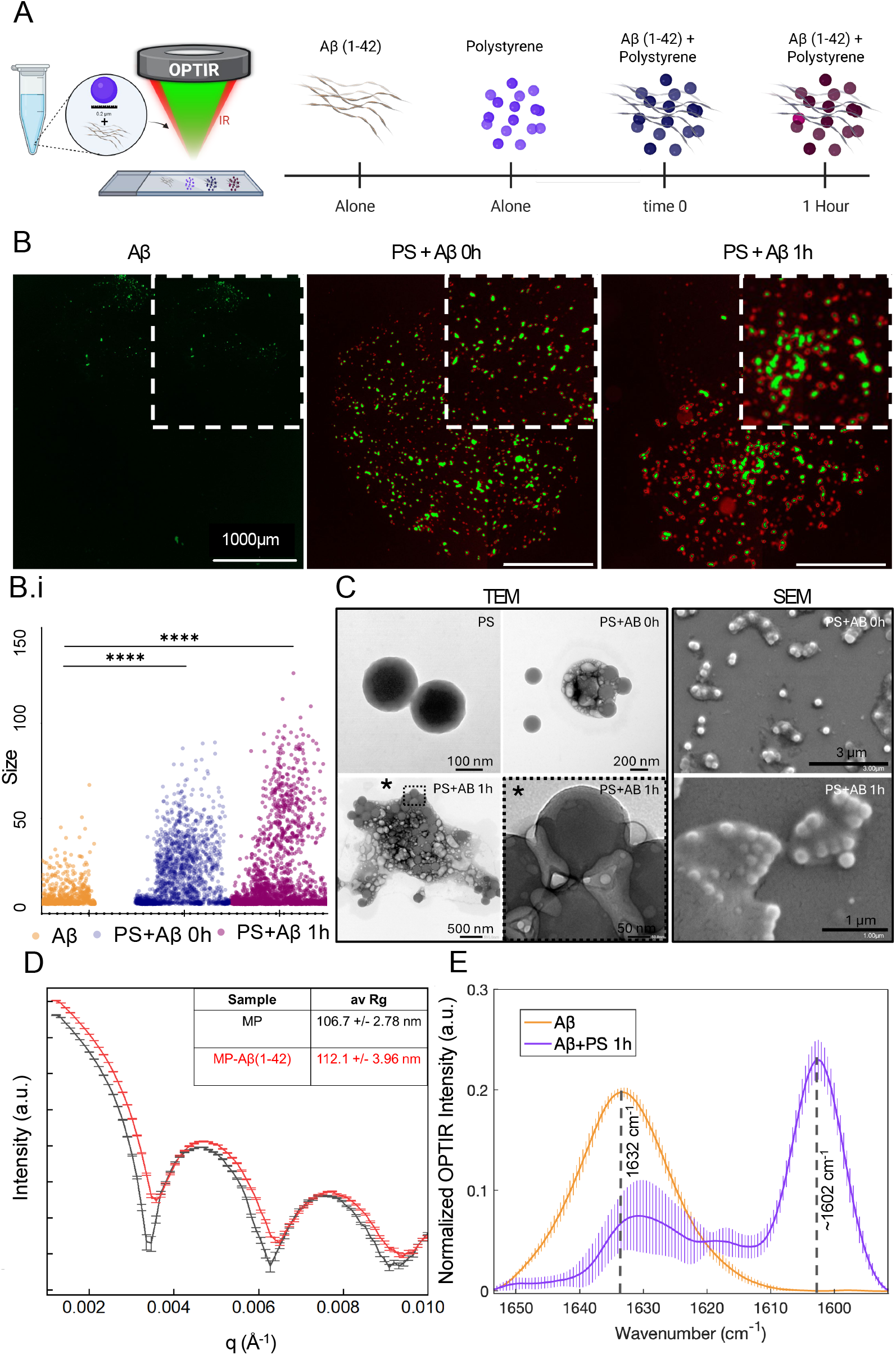
Characterization of the PS interaction with Aβ(1-42). **(A)** The experimental timeline is schematically created in BioRender. Silva, I. (2025) https://BioRender.com/sc8onrp, **(B)** Immunofluorescence imaging displays Aβ(1-42) in green and PS in red. (B.i) Adjacent to the images, a graph depicts the increase in aggregation from Aβ(1-42) alone to the PS+Aβ(1-42) combination at T0 and after one hour, (image scale, 1000 µm). **(C)** TEM/SEM representative images of the PS alone, Aβ(1-42) alone, to the PS + Aβ(1-42) at T0 and after one hour. **(D)** SAXS plot demonstrating changes in the average area of PS alone PS+Aβ combination. **(E)** O-PTIR spectra of Aβ(1-42) alone and PS + Aβ(1-42) after one hour. (Data representative of 3 independent experiments).

Further, corroborating the fluorescence microscopy data, Transmission Electron Microscopy (TEM) and Scanning Electron Microscopy (SEM) images in (**Figure 2C**) provide detailed ultrastructural insights. TEM images confirmed the approximate 200 nm size of individual PS. At T0 of the PS + Aβ (1-42) co-incubation, TEM revealed the early formation of aggregates, with distinct larger aggregate structures visible after 1 hour of co-incubation. SEM images provided even finer detail, showing how Aβ (1-42) protein appears to aggregate around the PS, forming larger, more complex structures. Final confirmation of aggregation and area increase was obtained from SAXS experiments (**Figure 2D**), which compared the average area of PS alone to that of the PS + Aβ (1-42) combination. This analysis demonstrated that the PS + Aβ (1-42) mixture formed significantly larger structures, with the radius of gyration of 112.1 ± 3.9 nm as compared to PS alone with the radius of gyration of106.7 ± 2.7. Finally, O-PTIR spectroscopy provided crucial molecular-level insights directly supporting Aβ1-42 fibrillization induced by the presence of PS. (**Figure 2E**) demonstrates O-PTIR spectra highlighting specific wavelengths characteristic of both PS and Aβ1-42, revealing the spectral interplay upon their co-incubation and indicating structural modifications due to their interaction. Specifically, the characteristic peak for PS at ~1600 cm^−1^ (also seen in **Figure 1**) and the Aβ (1-42) peak at ~1632 cm^−1^ are clearly observed in the same sample after 1 hour of co-exposure, confirming their co-presence and interaction during aggregation. This allowed us to compare amyloid structures formed by pure Aβ (1-42) and PS-bound Aβ (1-42). A notable strength of O-PTIR is its capacity to identify these interactions from the initial time point (0h) to 1h, enabling differentiation of individual materials at 0h and subsequent tracking the specific region for mapping or future analysis in the same area of their co-presence after exposure. The full spectral data, demonstrating the interrelation and progression across all four conditions (Aβ (1-42) alone, PS alone, Aβ (1-42) +PS at T0, and Aβ (1-42)+PS after 1 hour), can be seen in (**Supplementary Figure 1**).

### Dose-Dependent Cytotoxicity and Metabolic Impact of PS on Cells

Subsequently, we explored PS interaction in a more complex system. We conducted experiments on PS incorporated into N2Aswe cells, which increased the sample complexity while providing a well-defined and controlled environment. The experiments were performed with PS of a spherical shape, with an average diameter of 0.2 μm. Details on the PS and sample preparation are described in the methods section. We aimed to determine the optimal dose and exposure time for evaluating the effects of PS on cellular metabolism. To do this, we conducted two sets of experiments. Cells were analysed after 24 and 48 hours (**Supplementary Figure 2**). These findings help us evaluate the extent of changes and determine whether a 24-or 48-hours exposure is more appropriate for subsequent experiments.

To assess cellular uptake and cytotoxicity, we exposed cells to various concentrations of PS (0, 120, 240, and 480 µg/ml) for 24 and 48 hours. Bright-field microscopy (**Supplementary Figure 2 A**) revealed the uptake of PS by cells, visualized as yellow-highlighted regions indicating increased PS dose, and which also increases the fluorescence intensity.

To quantify the impact of PS on cell viability, we performed a WST-1 assay (**Supplementary Figure 2B**). We found a clear correlation between PS bead exposure and impaired cellular function. The WST-1 assay revealed a significant decrease in metabolic activity at all tested concentrations, suggesting PS induced cytotoxicity.

### Cytotoxicity Validation by LDH Assay

To definitively validate whether the observed metabolic impairment correlated with cellular damage or cell death, we performed the (LDH) cytotoxicity assay across the 24 hours exposure time-point (**Supplementary Figure 2C**). This assay directly measures LDH release into the medium, an indicator of plasma membrane integrity loss and irreversible cell death. The results confirm that the chosen PS concentrations did not induce high levels of cytotoxicity in either the N2Awt or N2Aswe cells.

Across all tested doses of PS 120 to 480 µg/mL, LDH release remained comparable to, or only slightly above, the untreated control groups, demonstrating that the doses selected for mechanistic study were sub-lethal and did not cause widespread cell death. Crucially, in N2Awt cells, neither the exposure to 10 mM Aβ(1-42) alone nor the co-exposure of 480µg/mL PS + 10 mM Aβ(1-42) resulted in acute cell death. This demonstrates that neither the Aβ(1-42) peptide alone nor its combination with the highest dose of PS is intrinsically cytotoxic at this early stage. In summary, while the WST-1 assay indicates a significant dose-dependent metabolic disturbance, the LDH assay confirms that this effect occurs without inducing severe cell death or overwhelming acute toxicity from the Aβ(1-42)/PS combination, validating the use of these doses for the study of structural and functional changes that precede apoptosis.

### Immunofluorescence-based analysis of cell viability loss, lysosomal engagement, and amyloid aggregation in response to PS exposure

To evaluate the impact of PS on cell viability and amyloid-related stress, we analysed immunofluorescence markers for lysosomes (Lamp1), apoptosis (Caspase-3), and Aβ different aggregation state (82e1, Aβ42, OC78, Amytracker) (**Figure 3A**), followed by (**Figure 3B**) providing an signal colocalization assessment plot of the interaction of the markers in the presence of PS.

**Figure 3.**
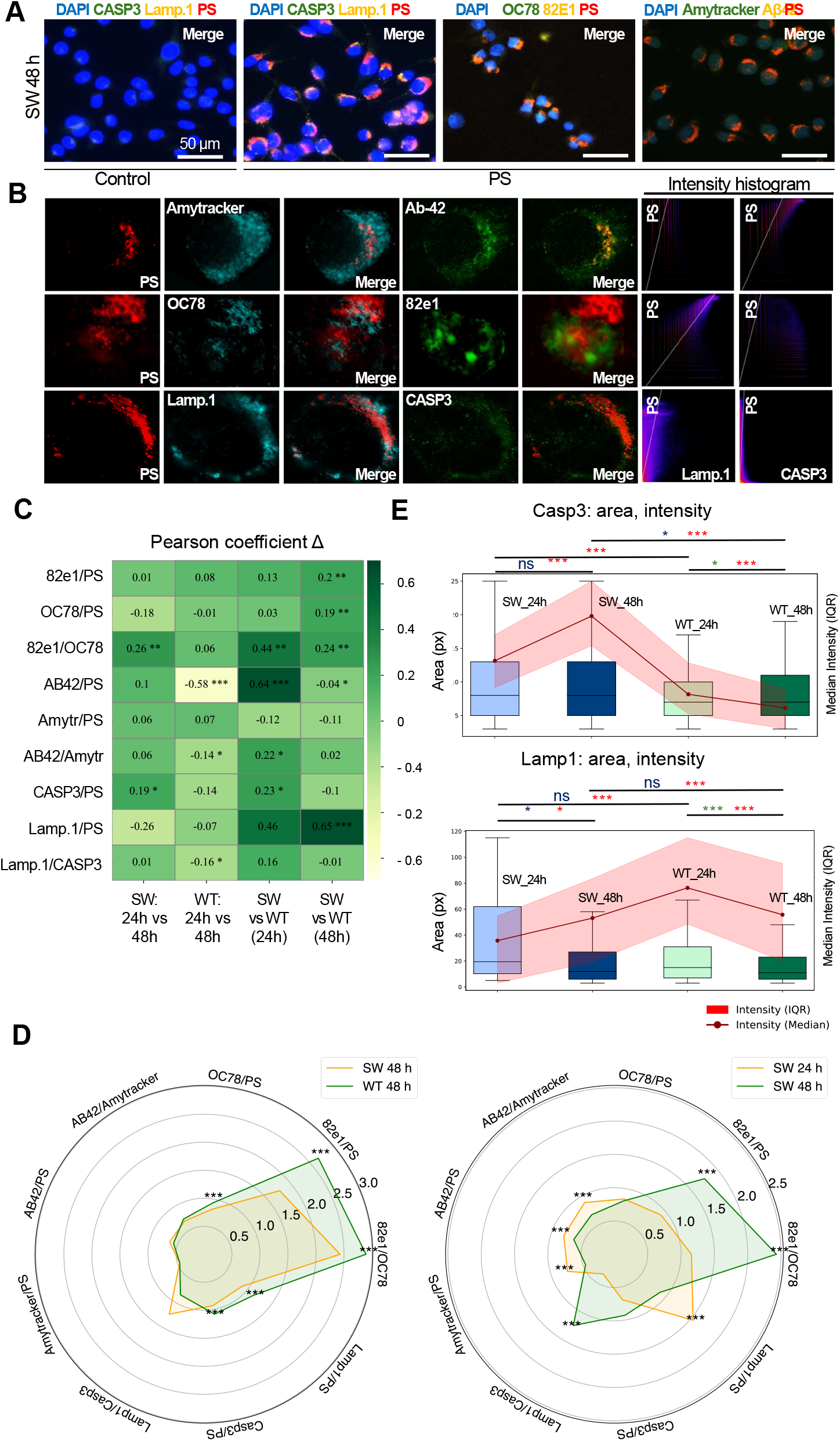
Cell viability loss, lysosomal engagement, and amyloid aggregation in response to PS exposure in N2Aswe (SW) and N2Awt (WT) cells at 24 h and 48 h. Scale bar: 50 μm. **(A)** Overview of the immunofluorescence panel used for the analysis of lysosomal (Lamp1), apoptotic (Caspase-3 (Casp3)), and Aβ (82e1, Aβ42, OC78, Amytracker) markers with PS particles. **(B)** Schematic workflow outlining the co-localization analysis process using Pearson and Manders metrics. **(C)** Pearson correlation coefficients quantifying the degree of spatial overlap between PS and target markers. **(D)** Manders M1/M2 ratio plots comparing co-localization trends across genotypes and timepoints. **(E)** 2D histograms showing distribution of positive signal area and intensity for Casp3 and Lamp1. (Data derived from n = 8 images per group. Error bars represent SEM).

### Lysosomal response and adaptation failure

At 24h, Lamp1intensity and area were elevated in both N2Awt (WT) and N2Aswe (SW) cells exposed to PS (**Figure 3E**), as well as in non-exposed controls (**Supplementary Figure 2A**), likely reflecting baseline activation after seeding. However, only PS-exposed WT and SW cells showed significant co-localization between Lamp1 and PS (Manders and Pearson, p < 0.05), confirming lysosomal uptake of PS particles (**Figure 3C and D**).

By 48h, Lamp1 levels in controls dropped significantly (p < 0.05), indicating normal adaptation in the absence of PS. In contrast, N2Awt and N2Aswe cells maintained high Lamp1 area and intensity, suggesting sustained lysosomal activity in response to PS exposure. Despite this, Lamp1–PS co-localization (Manders and Pearson) declined significantly in both groups from 24h to 48h (p < 0.05). The drop was sharper in N2Aswe, pointing to impaired lysosomal processing, likely due to the combined burden of PS and Aβ accumulation.

These results suggest that while both N2Awt and N2Aswe cells initially respond to PS via lysosomal engagement, N2Aswe cells fail to adapt by 48h, consistent with lysosomal dysfunction or degradation. The more pronounced Pearson decrease in N2Aswe supports the interpretation that PS–Aβ complexes are more resistant to clearance, increasing toxicity.

### Apoptosis is triggered by persistent PS and Aβ burden

Caspase-3 intensity increased over time in both genotypes but was significantly higher in N2Aswe at 48h (p < 0.05, **Figure 3E**). Co-localization with PS was present, but Manders M2 for Casp3– Lamp1 remained low, suggesting that apoptosis occurred near, but not within, lysosomal compartments (**Figure 3D**) (**Supplementary Figure 3C**). This supports the idea that PS aggregates not efficiently cleared by lysosomes initiate apoptotic stress, especially in N2Aswe cells, where Aβ burden exacerbates the effect. N2Awt cells also showed a statistically significant apoptotic response to PS, but to a lesser degree, confirming that PS alone is toxic, while the N2Aswe genotype amplifies the damage.

### PS enhances Aβ aggregation, accelerated in SW

At 24h, PS co-localized mainly with monomeric Aβ markers (82E1, Aβ42) in N2Aswe cells. By 48h, there was a shift toward increased labelling with OC78 which detects fibrillar oligomers and partially covers soluble oligomeric species and Amytracker, which specifically binds mature fibrils as indicated by elevated Manders M1 and Pearson coefficients (p < 0.05) (**Figure 3D**), indicating amyloid maturation around PS.

N2Awt cells statistically confirmed the trend of amyloid maturation, but with lower magnitude, suggesting that PS can nucleate Aβ aggregation, while N2Aswe accelerates the transition toward oligomeric and fibrillar forms associated with increased toxicity.

These findings strongly align with previous observations by [45], who demonstrated that nanoplastics can accelerate Aβ aggregation and potentiate neurotoxicity. PS likely act as catalytic surfaces for Aβ oligomerization via hydrophobic interactions, leading to increased burden on cellular clearance systems and loss of viability. These findings show that PS exposure induces lysosomal stress, amyloid aggregation, and apoptosis. In N2Aswe cells, the response is more sustained and dysfunctional, likely due to amyloid accumulation overwhelming degradation pathways. Even in N2Awt, prolonged PS exposure leads to cellular decline, supporting the hypothesis that PS nanoplastics are intrinsically toxic and exacerbate Aβ pathology when present.

### Structural Modifications of β-Sheets Induced by PS in vitro

Given that N2Aswe cells overexpress human APP and generate high levels of Aβ providing the necessary substrate to observe pathological misfolding, we focused our high-resolution O-PTIR structural analysis specifically on this AD model. To further investigate the effect of PS on protein structure at the subcellular level, we employed O-PTIR microscopy to assess β-sheet distribution in N2Aswe cells. Hyperspectral maps were analysed within the range of 1610 cm^−1^ – 1710 cm^−1^. First, as seen in (**Figure 4A**), we imaged the maps at specific wavenumbers: 1600 cm^−1^, indicative of PS-specific vibrations, and 1630 cm^−1^, associated with β-sheet structures. Both maps were normalized to 1658 cm^−1^, corresponding to total protein content (**Figure 4A**). The comparison of spectral maps from PS-exposed and unexposed N2Aswe cells corresponding to specific frequencies, revealed the intracellular presence and distribution of PS and β-sheet content. Notably, PS-exposed cells (N2Aswe+PS) exhibited a significant increase in the 1630 cm^−1^ signal compared (purple arrow) to control cells (grey arrow). These visualizations suggest elevated β-sheet levels in the presence of PS.

**Figure 4.**
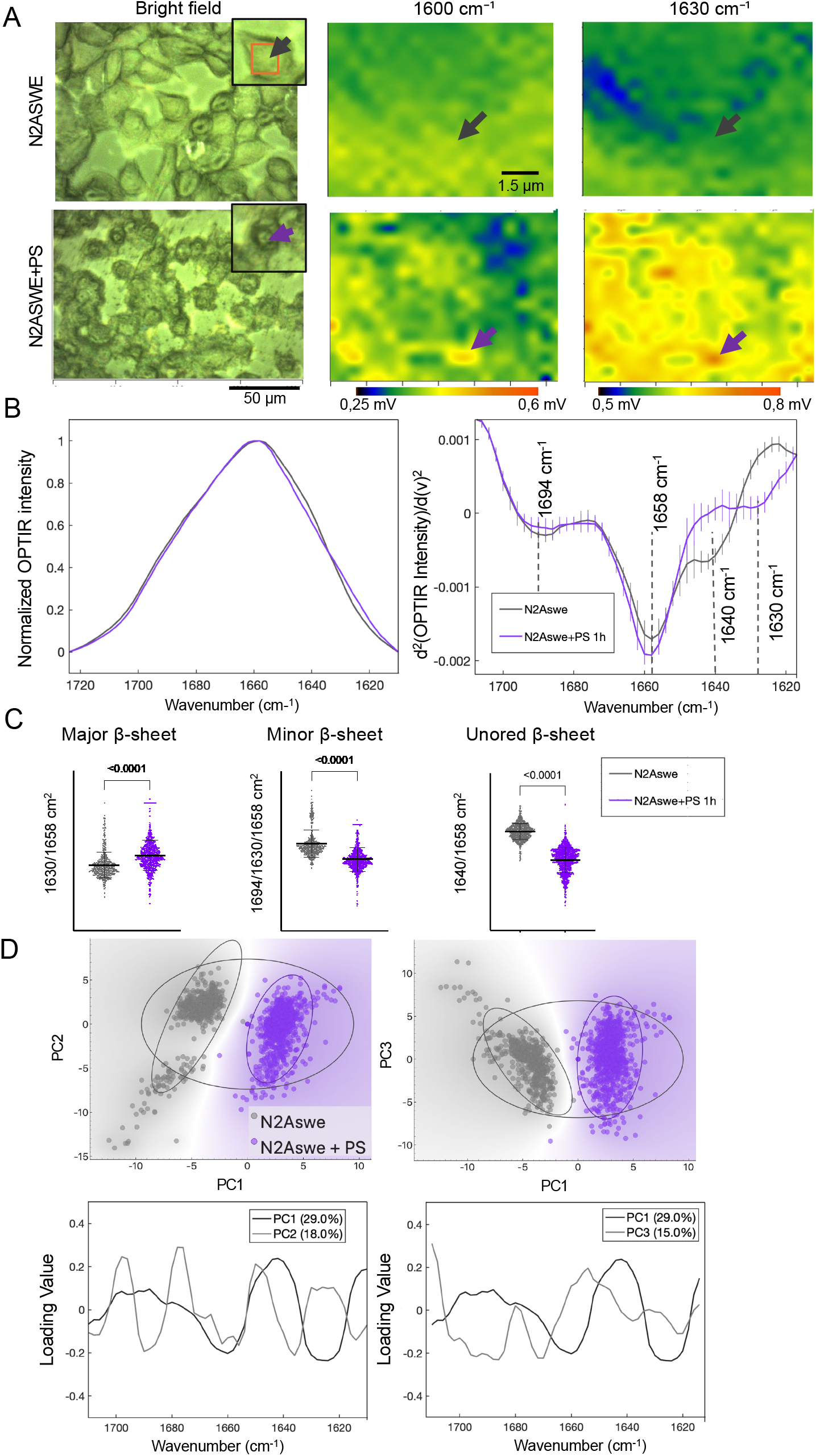
PS-induced changes in β-sheet structures in AD cellular model. **(A)** Brightfield image and hyperspectral O-PTIR maps of N2Aswe cells exposed or unexposed to PS, acquired at 1600cm^−1^ (PS-specific signal), 1630cm^−1^ (β-sheet content), normalized to 1658cm^−1^ (total protein/α-helices). PS-exposed cells showed increased intensity at 1630cm^−1^ (purple arrow), compared to non-exposed control (grey arrow). **(B)** Quantification of secondary structure components from second-derivative spectra of the Amide I region. Ratios were calculated between 1630cm^−1^ and 1658cm^−1^ (total β-sheets), 1694cm^−1^ and 1658/1630cm^−1^ (antiparallel β-sheets). **(C)** Principal Component Analysis (PCA) of Amide I spectra (1610–1710cm^−1^) showing clear separation between PS-exposed and control N2Aswe cells. PCA loading plots indicated that wavenumbers corresponding to β-sheet structures contributed most strongly to group differentiation. (Data derived from n = 5 (Control) and n = 6 (PS-exposed) individual cells, totaling >4,000 spectra. Statistical significance determined by Mann-Whitney U test).

To quantify the relationship between PS exposure and elevated β-sheet structures in N2Aswe, we calculated intensity ratios of secondary derivatives of Amide I band positions, which enhances the resolution of overlapping spectral components (**Figure 4B, Table 1**). This approach revealed that exposure to 480μg/mL PS for 24hours induced marked changes in the β-sheet content of N2Aswe cells. The total β-sheet level was assessed by calculating the intensity ratio of the 1630⍰cm^−1^ band relative to the 1658⍰cm^−1^ band. Antiparallel β-sheet structures, indicated by the 1694⍰cm^−1^ band, were quantified by their intensity ratios relative to both 1658⍰cm^−1^ and 1630⍰cm^−1^.

Likewise, unordered β-sheet structures at 1640⍰cm^−1^ were evaluated by calculating their intensity ratios relative to 1658cm^−1^. This approach normalizes β-sheet subtypes to both overall protein levels and total β-sheet content, providing a more detailed assessment of structural changes induced by PS exposure. This analysis confirmed that PS exposure results in an increase in β-sheet-rich content but decrease of antiparallel and unordered β-sheet structures, indicating a reorganization of protein conformation in response to PS exposure. According to [27], specific Amide I band positions reflect distinct β-sheet configurations and their hydrogen bonding environments.

The observed reduction in unordered and antiparallel β-sheets implies a shift toward more aggregated or fibrillar β-sheet forms. These structural rearrangements may compromise protein folding fidelity, potentially disrupting cellular homeostasis. Proper folding is critical for protein stability and function, and early conformational changes, such as those observed here, are key events in the pathogenesis of protein misfolding diseases like AD.

To ensure statistical robustness beyond the cell count, we analysed a comprehensive dataset of over 4,000 individual high-resolution spectra extracted from the hyperspectral maps. This massive spectral library allowed us to apply PCA to the spectra (**Figure 4C**). PCA is a multivariate statistical tool that reduces complex spectral data into orthogonal components (principal components), enabling the identification of wavenumbers that contribute most significantly to differences between groups [46]. This method facilitates the identification of underlying patterns and clustering based on spectral similarities. In our analysis, the first four components accounted for 70% of the total variance (**Figure 4C; Supplementary Figure 4**).

The PCA score plot clearly showed a distinct separation between PS-exposed and control in N2Aswe cell groups. It confirms that PS exposure induces systematic and quantifiable changes in the protein structure. The corresponding loading plots indicated that the wavenumbers contributing most strongly to group separation were those associated with β-sheet structural elements, further supporting our spectral interpretation.

Our findings indicate that exposure to PS can disrupt protein folding and stability by promoting β-sheet–rich conformations that are prone to aggregation. This structural shift may reflect an early step in pathological protein misfolding, which can affect cellular function. Given the wide presence of MNPs in the environment, it is crucial to determine whether similar protein-structural effects are triggered by other plastic types. Further investigation is needed to uncover the specific molecular mechanisms underlying PS interactions with β-sheet–containing proteins and to assess the long-term consequences of these interactions on human health.

Taken together, these data indicate that PS exposure contributes to intracellular protein aggregation, which consequently leads to metabolic dysregulation. These results align with studies demonstrating that MNPs can induce protein misfolding [47, 48] impair metabolic pathways [9], and reduce cell viability [49]. Such disruptions to proteostasis and energy balance are well-established drivers of disease progression in neurodegeneration, cardiovascular disorders, and cancer [6, 8, 50]. These disease-related processes are often driven by mechanisms such as increased oxidative stress, dysfunction of the protein quality control system, and long-term alterations in cellular metabolism [51–55]. Therefore, understanding how environmental contaminants like PS affect protein structure at the molecular level is crucial for a better understanding of microplastics potential role in AD and their impact on human health.

## DISCUSSION

The initial objective in understanding the biological impact of polystyrene (PS) microplastics was to assess the particle’s behaviour within the biological medium. We utilized commercially standardized 0.2 µm polystyrene beads (Sigma-Aldrich), a model widely established in nanotoxicology research. Previous physicochemical characterization of these specific particles in serum-supplemented media (DMEM with FBS) consistently reports the formation of a biomolecule corona. This is evidenced by a distinct zeta potential shift: while values in water are typically observed at approximately −45 mV, incubation in complete medium consistently shifts these values toward a stabilized range of −20 to −13 mV [56–59]. However, while Zeta potential infers surface interaction via charge alteration, we employed Small-Angle X-ray Scattering (SAXS) to directly quantify the physical dimensions of this complex. Our SAXS measurements (**Figure 2D**) confirmed a quantifiable increase in the radius of gyration (Rg) providing definitive structural verification of the Aβ coating in a physiological environment. This established profile indicates that serum proteins adsorb to the PS surface, providing steric stabilization that facilitates cellular uptake via endocytosis [57].

Our imaging data confirms this behaviour in our system, showing that PS particles remained dispersed as distinct puncta within the cytoplasm rather than forming large extracellular agglomerates. Consequently, the internalized PS surface acts as a reactive interface available for interaction with intracellular proteins, supporting our core hypothesis that the particle facilitates pathological protein processes [48, 58].

Our investigation demonstrated that the presence of PS microplastics significantly promotes amyloid protein aggregation. Through multi-modal analysis, including fluorescence microscopy, TEM, SEM, and SAXS, we observed that co-incubation of recombinant Aβ(1-42) with PS resulted in the formation of larger, more complex aggregated structures compared to Aβ(1-42) alone. O-PTIR spectroscopy provided critical molecular evidence, confirming the co-presence and interaction of PS (1600 cm^−1^) and Aβ(1-42) (1632 cm^−1^) within the same aggregate area. This evidence strongly supports the model that the PS microplastic acts as a nucleation scaffold or heterogeneous catalyst, accelerating the initial kinetic steps of Aβ aggregation [17]

While the in vitro data confirmed strong aggregation, rigorous environmental toxicology requires validation of the exposure dose. The WST-1 assay (**Supplementary Figure 2B**) indicated a significant, dose-dependent decrease in metabolic activity, suggesting initial cellular stress. To definitively validate whether this metabolic impairment progressed to irreversible cell death and to address reviewer concerns regarding the dosing strategy, we performed the LDH cytotoxicity assay (**Supplementary Figure 2C**). The results conclusively demonstrated that the PS concentrations used are sub-lethal in both N2Awt and N2Aswe cells at 24 hours, with LDH release remaining at basal levels. Crucially, in the N2Awt cells, neither 10 µM Aβ(1-42) alone nor the co-exposure of PS + Aβ(1-42) at the highest dose induced acute cell death. This validates our experimental approach: the doses are suitable for investigating sub-lethal mechanistic changes that precede acute toxicity [13, 52, 60].

The observed metabolic decline is underpinned by significant structural and cellular dysfunction. The co-localization of PS with the lysosomal marker Lamp1 confirms lysosomal uptake and engagement [59]. The failure to clear these aggregates, particularly in the N2Aswe cells, is indicated by the subsequent increase in Caspase-3, suggesting that the PS-Aβ complex initiates apoptotic signalling. The O-PTIR data provided the critical structural link. By focusing on the N2Aswe model, which offers a high density of amyloid targets, we were able to resolve that PS exposure induced marked structural alterations in cellular proteins, specifically evidenced by a significant increase in the 1630 cm^−1^ β- sheet signal and a reduction in antiparallel and unordered structures. This conformational rearrangement supports the hypothesis that the PS surface templates a shift toward highly aggregated, pathogenic Aβ forms. This structural shift, confirmed by Principal Component Analysis (PCA) clustering, validates that PS promotes pathological protein misfolding, which, combined with metabolic failure, drives toxicity [61]. We acknowledge that the PS concentrations utilized (120–480 µg/mL), while confirmed to be sub-lethal by our LDH analysis, are higher than current estimates of environmental exposure. However, these concentrations were specifically selected to align with the experimental framework established by Yang et al. (2022) [14] ensuring a comparable toxicological baseline. While their work identified that these doses induce ROS-mediated neuronal apoptosis, our study advances this finding by identifying the upstream molecular trigger: the surface-catalysed misfolding of amyloid proteins. The use of these established doses was technically requisite for O-PTIR analysis, the high signal-to-noise ratio allowed us to spectrally confirm the co-localization of PS and amyloid aggregates in situ a mechanistic confirmation that would be unresolvable at trace environmental levels. Consequently, this study serves as a mechanistic proof-of-concept; identifying the specific surface interactions that drive toxicity, which can now be targeted in future chronic exposure studies. Second, while the use of commercially standardized, pristine PS beads allows for reproducible physicochemical characterization, it does not fully capture the complexity of weathered, irregular microplastics found in nature, which may possess different surface chemistries and protein corona profiles. However, since the observed mechanism is driven by surface hydrophobicity and protein corona formation, we posit that this ‘catalytic scaffold’ effect is likely a shared feature across other hydrophobic polymers like polyethylene and polypropylene, broadening the potential environmental implications. Future work utilizing aged or environmentally sampled nanoplastics in complex co-culture models will be essential to further validate the toxicological pathway identified here.

## CONCLUSION

In conclusion, this study provides mechanistic evidence that exposure to polystyrene microplastics induces a dual cellular pathology in an in vitro model of Alzheimer’s Disease, within the promotion of pathological protein aggregation and the impairment of metabolic homeostasis. Utilizing O-PTIR spectroscopy, we identified specific structural alterations in cellular proteins, characterized by a distinct shift toward β-sheet-rich conformations and a reduction in unordered and antiparallel structures in N2Aswe cells. This conformational rearrangement, occurring alongside the co-localization of A-beta aggregates with the particles, strongly supports the hypothesis that the microplastic surface acts as a catalytic scaffold for amyloid nucleation.

Beyond structural misfolding, microplastic exposure disrupted metabolic function, evidenced by lysosomal stress and subsequent apoptotic signalling. Crucially, our LDH validation confirmed that these pathological changes occur at sub-lethal doses, distinguishing this mechanism from acute cytotoxicity and validating its relevance to chronic environmental toxicology. This work underscores the utility of O-PTIR as a powerful, high-resolution tool for investigating nanoscale molecular changes in response to environmental factors. Given the ubiquity of MNPs in the environment and their potential for bioaccumulation, this study identifies a plausible molecular pathway by which plastic pollution contributes to the risk and progression of neurodegenerative diseases, highlighting the urgent need for comprehensive human health risk assessments.

## LIST OF ABBREVIATIONS

MNPs: Microplastics and nanoplastics^1^
MPs: Microplastics (<5mm)^2^
NPs: Nanoplastics (<100 nm)^3^
PS: Polystyrene 4^4^
AD: Alzheimer’s Disease^5^
O-PTIR: Optical Photothermal Infrared Spectroscopy^6^
N2Awt: N2A wild type^7^
N2Aswe: N2A cell line stably transfected with human APP carrying the Swedish mutation^8^
DMEM: Dulbecco’s Modified Eagle Medium^9^
Opti-MEM: A cell culture medium^10^
FBS: Fetal Bovine Serum^11^
p/s: Penicillin/streptomycin^12^
CO2: Carbon dioxide^13^
WST-1: Water-soluble tetrazolium-1^14^
UV-vis: Ultraviolet-visible^15^
IR: Infrared^16^
QCL: Quantum cascade laser^17^
TEM: Transmission Electron Microscopy^18^
SEM: Scanning Electron Microscopy^19^
DAPI: 365/415 nm d ye^20^
CY5: 580/605 nm dye^21^
APP: Amyloid Precursor Protein^22^
Aβ: Amyloid beta^23^
HEWL: Hen egg-white lysozyme^24^
LPS: Lipopolysaccharide^25^
SAXS: Small Angle X-ray Scattering^26^
Rg: Radius of gyration^27^
WT: Wild type cells (N2Awt)^28^
SW: Swedish mutation cells (N2Aswe)^29^
PCA: Principal Component Analysis^30^

## AUTHOR INFORMATION

### Author Contributions

Iran Augusto Neves da Silva contributed to Conceptualization, Methodology, Formal Analysis, Investigation, Data Curation, Writing – Original Draft Preparation, Project Administration, and Funding Acquisition. Agnes Paulus contributed to Conceptualization, Methodology, Formal Analysis, Investigation, Data Curation, Writing – Review & Editing, and Project Administration. Valeriia Skoryk contributed to Methodology, Formal Analysis, Investigation, Data Curation, and Writing – Review & Editing. Kar-Yan Su contributed to Methodology, Investigation, and Writing Review & Editing., Herranz-Trillo, F contributed to Investigation and Writing – Review & Editing. Klementieva Oxana contributed to Conceptualization, Methodology, Investigation, Data Curation, Writing – Review & Editing, Project Administration, Resources, and Funding Acquisition. All authors have read and agreed to the published version of the manuscript.

## ACKNOWLEDGEMENT

We acknowledge the Microscopy for Biosciences (M-bio) Facility at the Faculty of Science, Department of Biology, Lund University. We also acknowledge the MAX IV Laboratory for beamtime on the CoSAXS beamline under proposal 20230450. Research conducted at MAX IV, a Swedish national user facility, is supported by the Swedish Research Council (grant number 2018-07152), Vinnova (grant number 2018-04969), Formas (grant number 2019-02496), Strategic Research Environment MultiPark and Nanolund (Multidisciplinary research on Parkinson’s disease), The Brain Foundation ( FO2022-0329) and The Royal Physiographic Society of Lund grant number F 2024/1892 and F 2024/1891. The Swedish Foundation for Strategic Research (grant number UKR24-0022). Crafoordska stiftelsen grant number (20250747). Olle Engkvists 220-0182 Forskningsprojekt.

**Data Availability The** raw O-PTIR spectral and SAXS data and imaging datasets generated during the current study are available from the corresponding author on reasonable request.

## CITATIONS

1. Jiang, B., et al., Health impacts of environmental contamination of micro-and nanoplastics: a review. Environmental health and preventive medicine, 2020. 25: p. 1–15.

2. Alimi, O.S., et al., Microplastics and Nanoplastics in Aquatic Environments: Aggregation, Deposition, and Enhanced Contaminant Transport. Environ Sci Technol, 2018. 52(4): p. 1704–1724.

3. Andrady, A.L. and M.A. Neal, Applications and societal benefits of plastics. Philosophical Transactions of the Royal Society B: Biological Sciences, 2009. 364(1526): p. 1977–1984.

4. Ebere, E.C., V.A. Wirnkor, and V.E. Ngozi, Uptake of microplastics by plant: a reason to worry or to be happy? World Scientific News, 2019(131):p. 256–267.

5. Enyoh, C.E., et al., Microplastics exposure routes and toxicity studies to ecosystems: an overview. Environmental Analysis, Health and Toxicology, 2020. 35(1).

6. Prata, J.C., et al., Environmental exposure to microplastics: An overview on possible human health effects. Science of the total environment, 2020. 702: p. 134455.

7. Hernandez, L.M., et al., Plastic teabags release billions of microparticles and nanoparticles into tea. Environmental science & technology, 2019. 53(21): p. 12300–12310.

8. Urli, S., et al., Impact of Microplastics and Nanoplastics on Livestock Health: An Emerging Risk for Reproductive Efficiency. Animals, 2023. 13(7): p. 1132.

9. Nihart, A.J., et al., Bioaccumulation of microplastics in decedent human brains. Nature Medicine, 2025:p. 1–6.

10. Bearer, E.L., et al., White matter hyperintensities and microplastics. bioRxiv, 2024:p. 2024.11.26.625277.

11. Chen, G.-f., et al., Amyloid beta: structure, biology and structure-based therapeutic development. Acta Pharmacologica Sinica, 2017. 38(9): p. 1205–1235.

12. Windheim, J., et al., Micro-and nanoplastics’ effects on protein folding and amyloidosis. International journal of molecular sciences, 2022. 23(18): p. 10329.

13. Shan, S., et al., Polystyrene nanoplastics penetrate across the blood-brain barrier and induce activation of microglia in the brain of mice. Chemosphere, 2022. 298: p. 134261.

14. Yang, D., et al., Polystyrene micro- and nano-particle coexposure injures fetal thalamus by inducing ROS-mediated cell apoptosis. Environ Int, 2022. 166: p. 107362.

15. Mahmoudi, M., et al., Protein fibrillation and nanoparticle interactions: opportunities and challenges. Nanoscale, 2013. 5(7): p. 2570–2588.

16. Hensley, K., et al., A model for beta-amyloid aggregation and neurotoxicity based on free radical generation by the peptide: relevance to Alzheimer disease. Proceedings of the National Academy of Sciences, 1994. 91(8): p. 3270–3274.

17. Cabaleiro-Lago, C., et al., Dual effect of amino modified polystyrene nanoparticles on amyloid β protein fibrillation. ACS chemical neuroscience, 2010. 1(4): p. 279–287.

18. Salminen, A., A. Kauppinen, and K. Kaarniranta, Emerging role of NF-κB signaling in the induction of senescence-associated secretory phenotype (SASP). Cellular signalling, 2012. 24(4): p. 835–845.

19. Varela-Eirín, M. and M. Demaria, Cellular senescence. Current Biology, 2022. 32(10): p. R448–R452.

20. Yang, D., et al., Polystyrene micro-and nano-particle coexposure injures fetal thalamus by inducing ROS-mediated cell apoptosis. Environment International, 2022. 166: p. 107362.

21. Yang, Q., et al., Oral feeding of nanoplastics affects brain function of mice by inducing macrophage IL-1 signal in the intestine. Cell Reports, 2023. 42(4).

22. Selkoe, D.J., Alzheimer’s disease results from the cerebral accumulation and cytotoxicity of amyloid ß-protein. Journal of Alzheimer’s disease, 2001. 3(1): p. 75–80.

23. Gouras, G.K., T.T. Olsson, and O. Hansson, β-Amyloid peptides and amyloid plaques in Alzheimer’s disease. Neurotherapeutics, 2015. 12(1): p. 3–11.

24. Gvazava, N., et al., Label-free high-resolution photothermal optical infrared spectroscopy for spatiotemporal chemical analysis in fresh, hydrated living tissues and embryos. Journal of the American Chemical Society, 2023. 145(45): p. 24796–24808.

25. Prater, C.B., M. Kansiz, and J.-X. Cheng, A tutorial on optical photothermal infrared (O-PTIR) microscopy. APL Photonics, 2024. 9(9).

26. Cheng, J.-X. and X.S. Xie, Coherent anti-Stokes Raman scattering microscopy: instrumentation, theory, and applications. 2004, ACS Publications. p. 827–840.

27. Barth, A., Infrared spectroscopy of proteins. Biochimica et Biophysica Acta (BBA)-Bioenergetics, 2007. 1767(9): p. 1073–1101.

28. Cerf, E., et al., Antiparallel β-sheet: a signature structure of the oligomeric amyloid β-peptide. Biochemical Journal, 2009. 421(3): p. 415–423.

29. Thinakaran, G., et al., Metabolism of the “Swedish” amyloid precursor protein variant in neuro2a (N2a) cells: evidence that cleavage at the “β-secretase” site occurs in the Golgi apparatus (*). Journal of Biological Chemistry, 1996. 271(16): p. 9390–9397.

30. Da Silva, I.A., et al., Formalin-free fixation and xylene-free tissue processing preserves cell-hydrogel interactions for histological evaluation of 3D calcium alginate tissue engineered constructs. Frontiers in Biomaterials Science, 2023. 2: p. 1155919.

31. Ami, D., P. Mereghetti, and A. Natalello, Contribution of infrared spectroscopy to the understanding of amyloid protein aggregation in complex systems. Frontiers in Molecular Biosciences, 2022. 9: p. 822852.

32. Geraldes, C.F., Introduction to infrared and Raman-based biomedical molecular imaging and comparison with other modalities. Molecules, 2020. 25(23): p. 5547.

33. Kansiz, M. and C. Marcott. Submicron simultaneous IR and Raman microscopy (IR+ Raman): breakthrough developments in Optical Photothermal IR (O-PTIR) combined with Raman provide new capabilities. in AOS Australian Conference on Optical Fibre Technology (ACOFT) and Australian Conference on Optics, Lasers, and Spectroscopy (ACOLS) 2019. 2019. SPIE.

34. Kansiz, M., et al., Optical photothermal infrared microspectroscopy with simultaneous Raman–a new non-contact failure analysis technique for identification of< 10 μm organic contamination in the hard drive and other electronics industries. Microscopy today, 2020. 28(3): p. 26–36.

35. Zhang, D., et al., Depth-resolved mid-infrared photothermal imaging of living cells and organisms with submicrometer spatial resolution. Science advances, 2016. 2(9): p. e1600521.

36. Demšar, J. and B. Zupan, Hands-on training about data clustering with orange data mining toolbox. PLOS Computational Biology, 2024. 20(12): p. e1012574.

37. Confer, M.P., et al., Label-free infrared spectroscopic imaging reveals heterogeneity of β-sheet aggregates in Alzheimer’s disease. The journal of physical chemistry letters, 2021. 12(39): p. 9662–9671.

38. Holcombe, B., et al., Intermediate Antiparallel β Structure in Amyloid β Plaques Revealed by Infrared Spectroscopic Imaging. ACS chemical neuroscience, 2023. 14(20): p. 3794–3803.

39. Ganim, Z., et al., Amide I two-dimensional infrared spectroscopy of proteins. Accounts of chemical research, 2008. 41(3): p. 432–441.

40. Duswald, K., et al., Detection of Unlabeled Micro-and Nanoplastics in Unstained Tissue with Optical Photothermal Infrared Spectroscopy. bioRxiv, 2024:p. 2024.11.11.622943.

41. Petoukhov, M.V., et al., ATSAS 2.1–towards automated and web-supported small-angle scattering data analysis. Applied crystallography, 2007. 40(S1): p. s223–s228.

42. Manalastas-Cantos, K., et al., ATSAS 3.0: expanded functionality and new tools for small-angle scattering data analysis. Applied Crystallography, 2021. 54(1): p. 343–355.

43. Painter, P., M. Sobkowiak, and Y. Park, Vibrational relaxation in atactic polystyrene: an infrared spectroscopic study. Macromolecules, 2007. 40(5): p. 1730–1737.

44. Chatzi, E.G., et al., Infrared spectra and compositional analysis of styrene/2-ethylhexyl acrylate copolymers. Macromolecular Chemistry and Physics, 1997. 198(8): p. 2409–2420.

45. Gou, X., et al., Impact of nanoplastics on Alzheimer’s disease: Enhanced amyloid-β peptide aggregation and augmented neurotoxicity. Journal of Hazardous Materials, 2024. 465: p. 133518.

46. Groth, D., et al., Principal components analysis. Computational Toxicology: Volume II, 2013:p. 527–547.

47. Bearer, E.L., et al., White matter hyperintensities and microplastics. bioRxiv, 2024:p. 2024.11.26.625277.

48. Chen, Y., et al., The impact of modified polystyrene on lysozyme fibrillation studied by surface-enhanced Raman spectroscopy (SERS). International Journal of Biological Macromolecules, 2023. 242: p. 124937.

49. Murali, K., et al., Uptake and bio-reactivity of polystyrene nanoparticles is affected by surface modifications, ageing and LPS adsorption: in vitro studies on neural tissue cells. Nanoscale, 2015. 7(9): p. 4199–4210.

50. Rist, S., et al., A critical perspective on early communications concerning human health aspects of microplastics. Science of the Total Environment, 2018. 626: p. 720–726.

51. Lu, L., et al., Polystyrene microplastics induce gut microbiota dysbiosis and hepatic lipid metabolism disorder in mice. Science of the total environment, 2018. 631: p. 449–458.

52. Hu, M. and D. Palić, Micro-and nano-plastics activation of oxidative and inflammatory adverse outcome pathways. Redox biology, 2020. 37: p. 101620.

53. Gopinath, P.M., et al., Prospects on the nano-plastic particles internalization and induction of cellular response in human keratinocytes. Particle and Fibre Toxicology, 2021. 18(1): p. 1–24.

54. Ehsanifar, M., et al., Hippocampal inflammation and oxidative stress following exposure to diesel exhaust nanoparticles in male and female mice. Neurochemistry International, 2021. 145: p. 104989.

55. Zhang, H., et al., Combined exposure of alumina nanoparticles and chronic stress exacerbates hippocampal neuronal ferroptosis via activating IFN-γ/ASK1/JNK signaling pathway in rats. Journal of Hazardous Materials, 2021. 411: p. 125179.

56. Hou, Y., et al., An integrative method for evaluating the biological effects of nanoparticle-protein corona. Biochimica et Biophysica Acta (BBA) - General Subjects, 2023. 1867(3): p. 130300.

57. Han, S., et al., Endosomal sorting results in a selective separation of the protein corona from nanoparticles. Nature Communications, 2023. 14(1): p. 295.

58. Tallec, K., et al., Surface functionalization determines behavior of nanoplastic solutions in model aquatic environments. Chemosphere, 2019. 225: p. 639–646.

59. Mota, C., et al., Are all nanoplastics equally neurotoxic? Influence of size and surface functionalization on the toxicity of polystyrene nanoplastics in human neuronal cells. Environmental Pollution, 2026. 390: p. 127445.

60. Ghosal, S., S. Bag, and S. Bhowmik, Insights into the binding interactions between microplastics and human α-synuclein protein by multispectroscopic investigations and amyloidogenic oligomer formation. The Journal of Physical Chemistry Letters, 2024. 15(25): p. 6560–6567.

61. Klementieva, O., et al., Super-Resolution Infrared Imaging of Polymorphic Amyloid Aggregates Directly in Neurons. Advanced Science, 2020. 7(6): p. 1903004.

