## Supplemental figures for "Microplastic Exposure Promotes Amyloid Misfolding and Metabolic Impairment at Sub-Lethal Doses in an In Vitro Cellular Model of Alzheimer’s Disease"

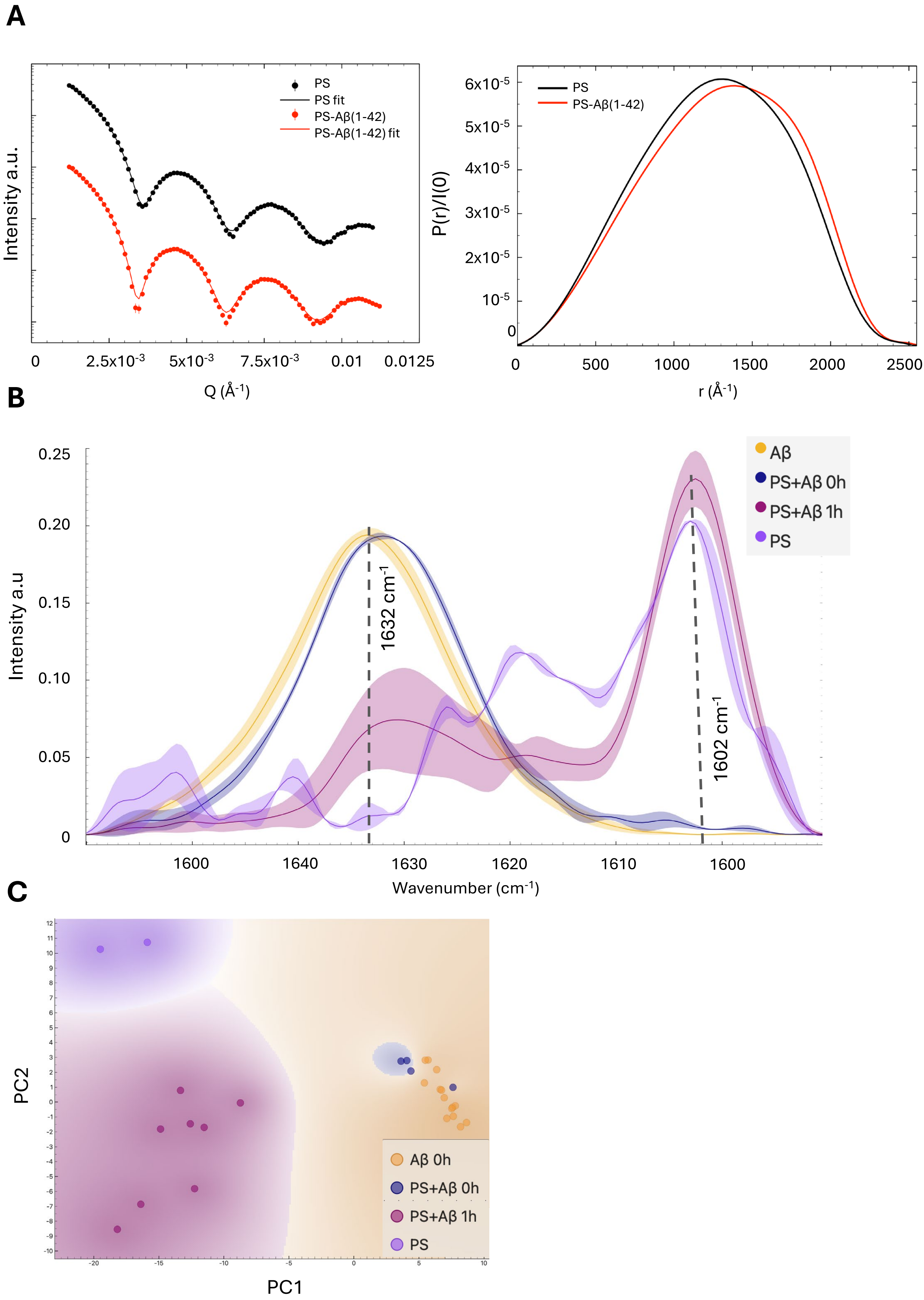


**Supplementary Figure S1: Comprehensive O-PTIR Spectral Analysis and PCA of Aβ (1-42) and PS Interaction. A.** Pair distance distribution, p(r), profile normalized with I(0) (right) and fit with experimental data (left). The difference in p(r) shape indicate a slightly more elongate shape of the complex compared with the PS alone. **B.** O-PTIR spectra showing the interrelation and progression across all four conditions: **Aβ (1-42)** alone, PS alone, **Aβ (1-42)** +PS at time zero (T0), and **Aβ (1-42)** +PS after 1 hour of co−incubation. The spectra illustrate the distinct absorption peaks for PS (around 1602cm −1 ) and **Aβ (1-42)** (around ~1632 $cm^{-1}$), demonstrating their co-presence and spectral interplay upon interaction. The integrity of the **Aβ (1-42)** protein was confirmed by the presence of the characteristic Amide I band around 1633 $cm^{-1}$, serving as an internal control for any conformational changes. **C.** Principal Component Analysis (PCA) scatter plot illustrating the spectral variations and grouping of samples from PS alone, **Aβ (1-42)** alone, and their mixture at T0 and after 1 hour. This analysis visually confirms the distinct spectral profiles of the individual components and the changes occurring upon co-incubation.

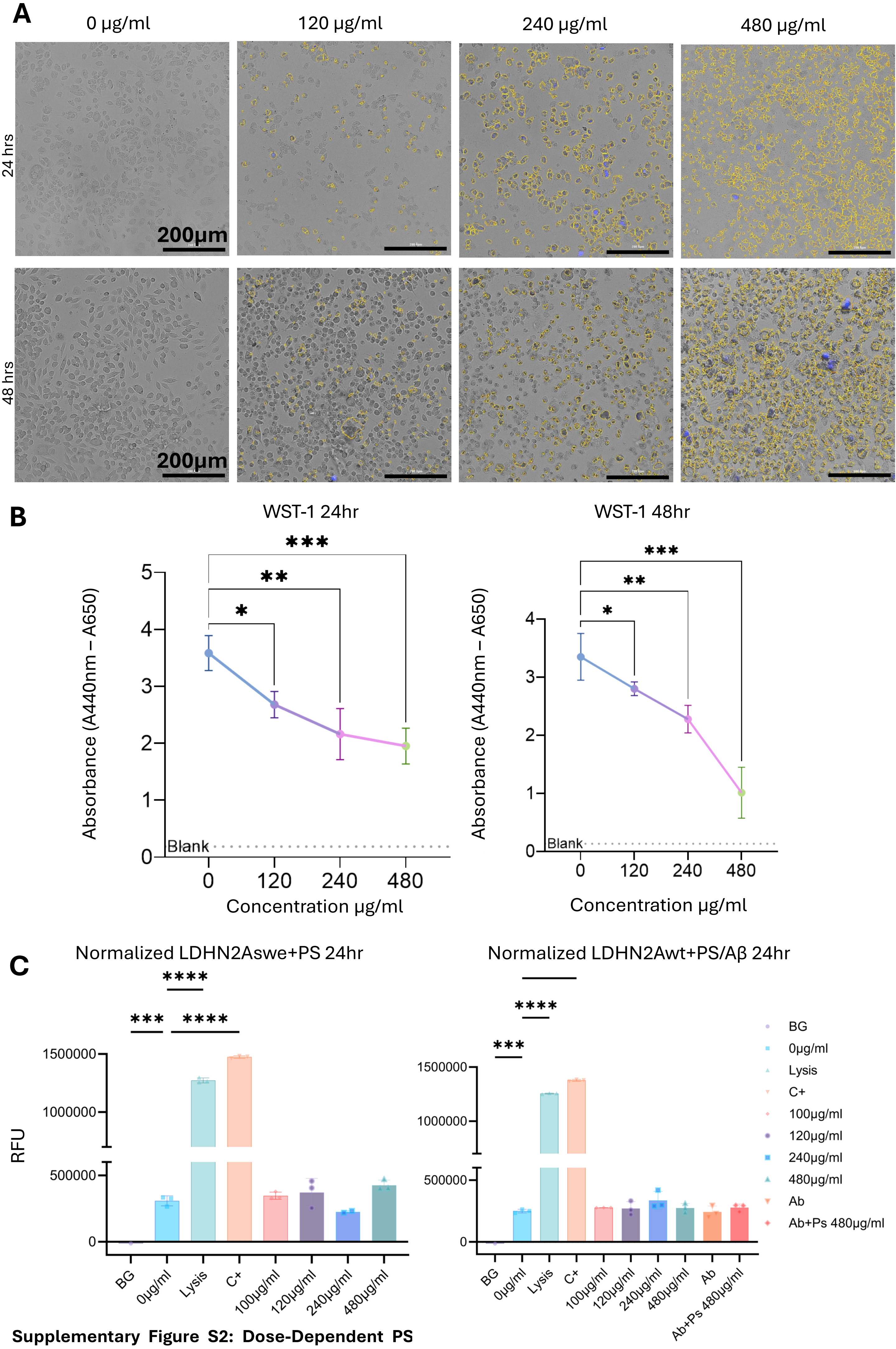


**Supplementary Figure S2: Dose-Dependent PS Viability. A.** Bright-field microscopy images of N2Aswe cells, illustrating the effects of increasing concentrations of polystyrene (PS) (0 µg/ml, 120 µg/ml, 240 µg/ml, and 480 µg/ml) after 24- and 48-hour exposure periods. Yellow-highlighted regions indicate the intracellular uptake and accumulation of PS, which increases with higher PS doses. Scale bar, 200 µm. **B.** Graph of Water-Soluble Tetrazolium-1 (WST-1) assay results, demonstrating the dose-dependent decrease in N2Aswe cellular metabolic activity upon exposure to increasing concentrations of PS. This indicates PS-induced cytotoxicity. **C.** Quantification of LDH release in N2Awt and N2Aswe cells. The results confirm that all tested doses of PS alone (120 to 480 µg/ml) induce sub-acute cytotoxicity, with LDH release remaining low and comparable to controls. Crucially, in N2Awt cells, neither the exposure to 10 mM Aβ(1-42) alone nor the co-exposure of 480 µg/ml PS + 10 mM Aβ(1-42) resulted in significant acute cell death.

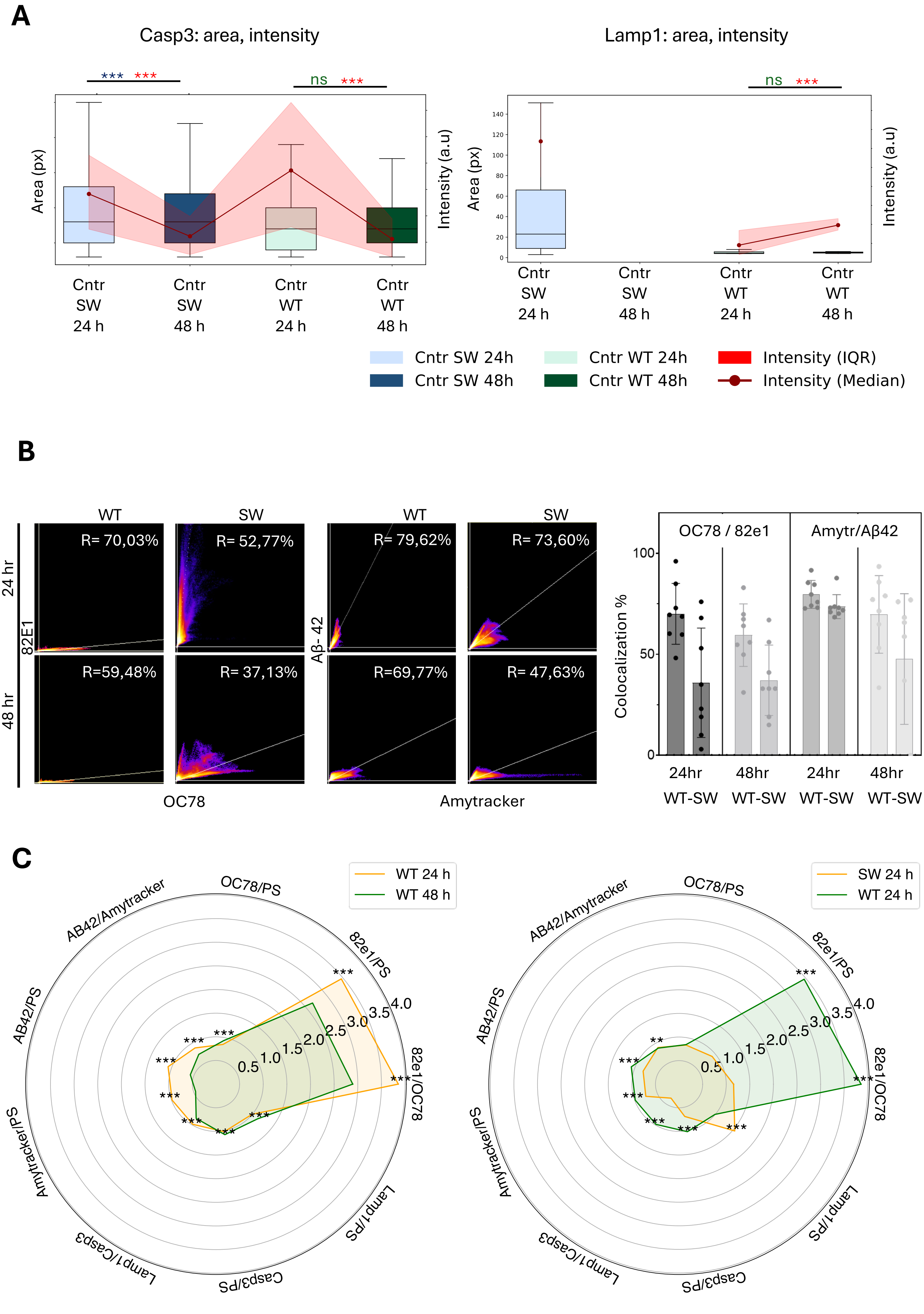


**Supplementary Figure S3: Baseline Levels of Apoptotic and Lysosomal Markers in Control Cells, and Detailed Co-localization Trends with PS.** A. Box plots illustrating the baseline levels of Caspase-3 (Casp3) area and intensity in control (non-exposed) N2Aswe (SW) and N2Awt (WT) cells at 24h and 48h. This panel provides a baseline for comparison with PS-exposed cells. B. Box plots illustrating the baseline levels of LAMP1 area and intensity in control (non-exposed) N2Aswe (SW) and N2Awt (WT) cells at 24h and 48h. This panel shows the natural variability and adaptation of lysosomal activity in control conditions over time. C. Radar plots providing a comprehensive overview of the co-localization trends between PS and various cellular markers (82e1, OC78, Aβ42, Amytracker for amyloid states; LAMP1 for lysosomes; Caspase-3 for apoptosis) in N2Aswe (SW) and N2Awt (WT) cells at 24h and 48h. These plots offer an alternative visualization of the co-localization data presented in main Figure 3D, highlighting the relative changes in association between PS and these markers across different genotypes and exposure durations.

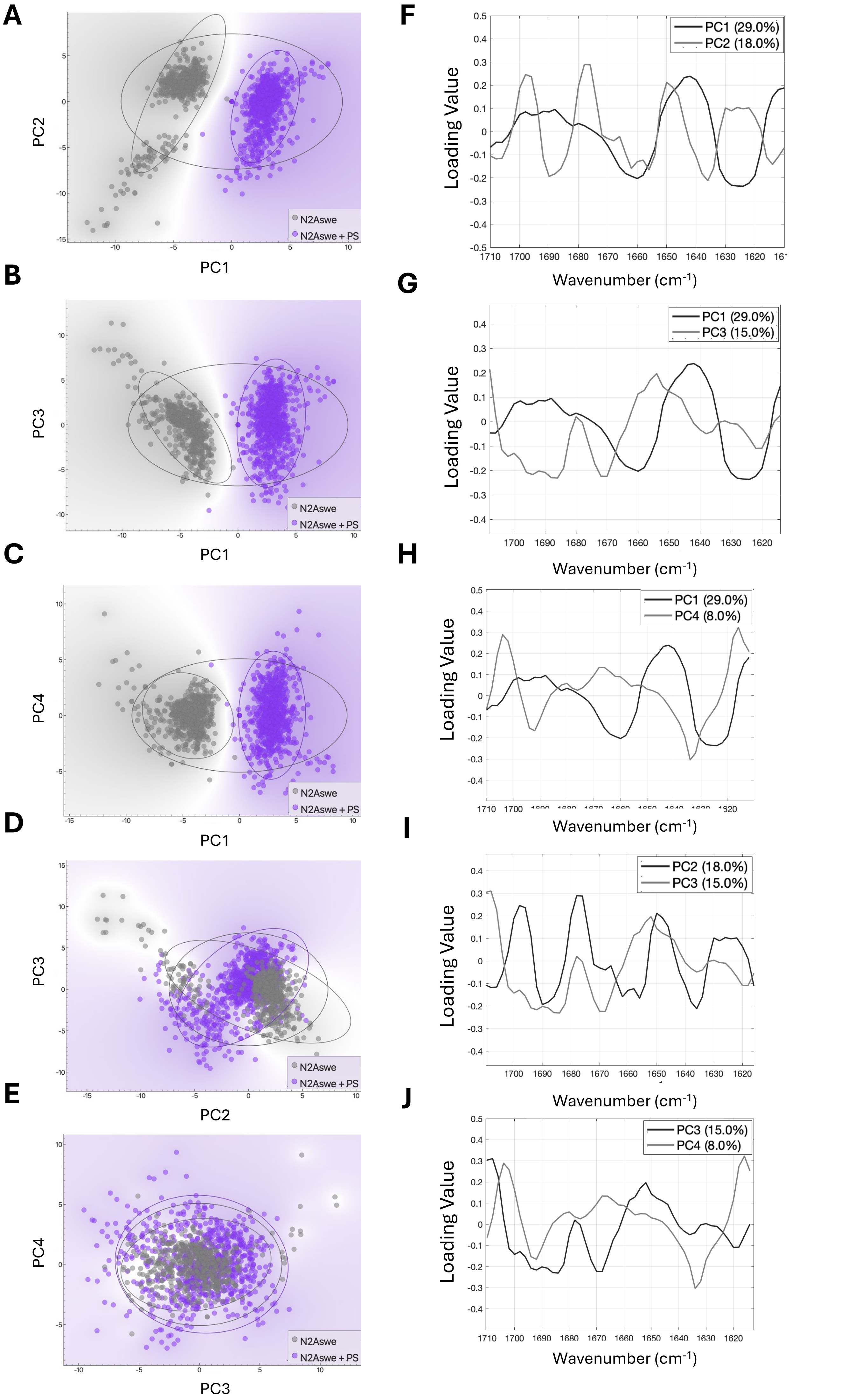


**Supplementary Figure S4: Principal Component Analysis (PCA) of O-PTIR Spectra from N2Aswe Cells with and Without PS Exposure.** A-E. PCA scatter plots (showing various combinations of PC1, PC2, PC3, and PC4) illustrating the clear separation between PS-exposed (purple) and control (grey) N2Aswe cell groups based on their O-PTIR spectral profiles in the Amide I region (1610–1710 $cm^{-1}$). This confirms that PS exposure induces systematic and quantifiable changes in cellular protein structure. F-J. Corresponding PCA loading plots for PC1, PC2, PC3, and PC4, indicating the specific wavenumbers that contribute most strongly to the observed group separation. These plots further support that differences are primarily associated with β-sheet structural elements (e.g., around 1630 $cm^{-1}$, 1658 $cm^{-1}$, 1694 $cm^{-1}$).
